# Responses to song playback differ in sleeping versus anesthetized songbirds

**DOI:** 10.1101/2022.01.11.475714

**Authors:** Sarah W. Bottjer, Chloé Le Moing, Ellysia Li, Rachel Yuan

## Abstract

Vocal learning in songbirds is mediated by a highly localized system of interconnected forebrain regions, including recurrent loops that traverse the cortex, basal ganglia, and thalamus. This brain-behavior system provides a powerful model for elucidating mechanisms of vocal learning, with implications for learning speech in human infants, as well as for advancing our understanding of skill learning in general. A long history of experiments in this area has tested neural responses to playback of different song stimuli in anesthetized birds at different stages of vocal development. These studies have demonstrated selectivity for different song types that provide neural signatures of learning. In contrast to the ease of obtaining responses to song playback in anesthetized birds, song-evoked responses in awake birds are greatly reduced or absent, indicating that behavioral state is an important determinant of neural responsivity. Song-evoked responses can be elicited in sleeping as well as anesthetized zebra finches, and the selectivity of responses to song playback in adult birds tends to be highly similar between anesthetized and sleeping states, encouraging the idea that anesthesia and sleep are highly similar. In contrast to that idea, we report evidence that cortical responses to song playback in juvenile zebra finches (*Taeniopygia guttata*) differ greatly between sleep and urethane anesthesia. This finding indicates that behavioral states differ in sleep versus anesthesia and raises questions about relationships between developmental changes in sleep activity, selectivity for different song types, and the neural substrate for vocal learning.

**Significance:** Patterns of spiking activity based on electrophysiological recordings in many different taxa are known to be heavily dependent on behavioral state. Neural activity patterns are frequently similar between sleep and anesthesia, which has encouraged the idea that similar states characterize sleep and anesthesia. Based on comparisons across studies, we report that activity patterns are highly dissimilar between sleep and urethane anesthesia in a cortical region of juvenile songbirds. These data argue against the idea that similar behavioral states are achieved in sleep versus anesthesia.

## Introduction

Vocal learning in zebra finches serves as a powerful model for investigating mechanisms of motor skill learning during development (Doupe and Kuhl, 1999; Brainard and Doupe, 2013). Juvenile zebra finches learn the sounds used for vocal communication, and this type of skill learning, like other forms of goal-directed learning, is controlled by cortico-basal ganglia circuits (Yin and Knowlton, 2006; Graybiel, 2008; Redgrave et al., 2010; Turner and Desmurget, 2010; Cox and Witten, 2019). Similar to infants learning speech, juvenile songbirds memorize the vocal sounds of their adult tutor. They then progressively refine their own vocal behavior to imitate the tutor song (the goal behavior) during the sensorimotor stage of vocal learning. This process requires the evaluation of feedback of self-generated vocalizations against a neural representation of the goal tutor song to guide the gradual acquisition of an accurate imitation.

Neural control of vocal learning in juvenile zebra finches is vested in basal ganglia loops that emanate from the cortical nucleus LMAN (Fig. 1) (Bottjer et al., 1984; Scharff and Nottebohm, 1991; Aronov et al., 2008). CORE and SHELL subregions of LMAN make parallel connections through the basal ganglia and thalamus (Johnson et al., 1995; Iyengar et al., 1999; Luo et al., 2001; Iyengar and Bottjer, 2002b; Bottjer, 2004; Gale et al., 2008; Person et al., 2008; Paterson and Bottjer, 2017). The CORE pathway mediates vocal motor production in juvenile songbirds (Bottjer et al., 1984; Scharff and Nottebohm, 1991; Aronov et al., 2008; Elliott et al., 2014; Kojima et al., 2018) and is functionally similar to sensorimotor cortico-basal ganglia loops in mammals that contribute to learning and performance (Alexander and Crutcher, 1990; Graybiel, 2008; Yin et al., 2009; Ashby et al., 2010; Redgrave et al., 2010; Thorn et al., 2010; Gremel and Costa, 2013; Kupferschmidt et al., 2017). In contrast, the SHELL pathway is involved in evaluating sensorimotor performance and is functionally similar to associative-limbic loops that traverse the basal ganglia; lesions in the SHELL pathway of juvenile birds impair the ability to imitate tutor song, but do not cause motor disruption of song production (Bottjer and Altenau, 2010). This disruption of learning but not motor performance suggests that SHELL circuitry may help to evaluate whether self-generated vocalizations match learned tutor sounds.

**Figure 1.**
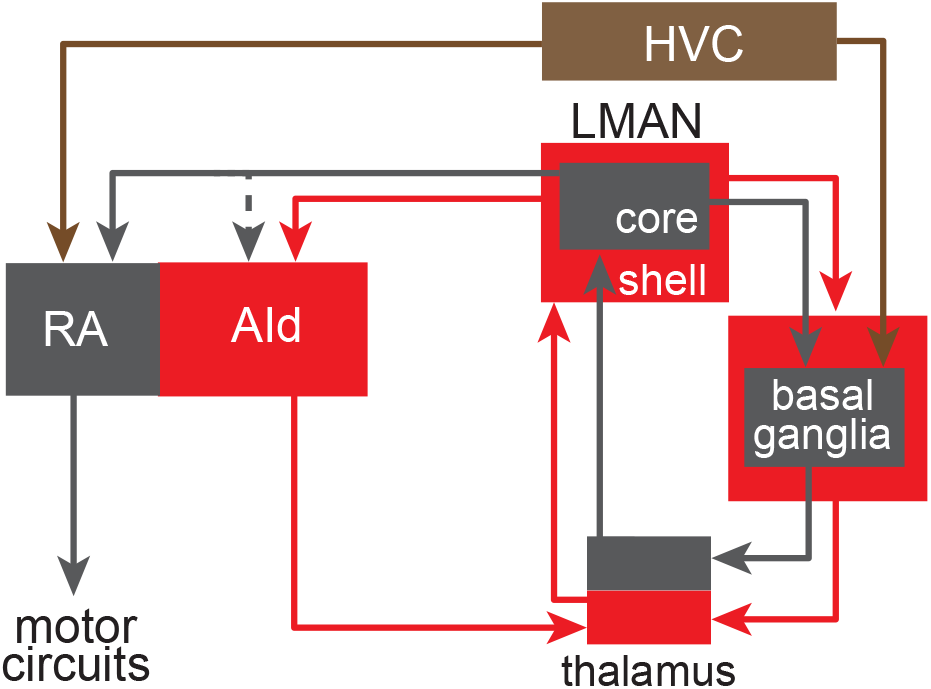
A simplified schematic of cortico-basal ganglia circuits that mediate vocal learning and behavior. The cortical nucleus LMAN comprises CORE (gray) and SHELL (red) subregions which form parallel recurrent loops through the basal ganglia and dorsal thalamus. LMAN-SHELL also forms a trans-cortical loop via AId that converges with basal ganglia loops in the same dorsal thalamic zone. A transient projection from LMAN-CORE to AId is present only in juvenile birds and creates a site of integration between CORE and SHELL pathways in AId during early sensorimotor learning (denoted by dotted line). The dorsal thalamic zone feeds back to LMAN and feeds forward to HVC via medial MAN (latter pathway not shown for clarity). A specific region of the basal ganglia known as Area X is dedicated to functions for vocal learning and includes both striatal and pallidal cells. Abbreviations: RA: robust nucleus of the arcopallium; AId: dorsal intermediate arcopallium; HVC: high vocal center; LMAN: lateral magnocellular nucleus of the anterior nidopallium.

Studies of the mechanisms that underlie vocal learning in songbirds have a long and venerable history of examining neural responses to playback of different song types in anesthetized juvenile and adult birds (Margoliash, 1983, 1986; Volman, 1993; Lewicki and Konishi, 1995; Solis and Doupe, 1997, 1999, 2000; Adret et al., 2012). One quest in this area was to discover a population of neurons that encode the tutor song memorized by each juvenile bird. Achiro and Bottjer (2013) reported that the SHELL subregion of LMAN in juvenile anesthetized birds contains a large proportion of neurons (^~^30%) that respond significantly only to playback of tutor song. This tutor-tuned population provides a target memory that is essential for matching self-generated utterances to the goal tutor song, and is present only during early stages of sensorimotor integration. The proportion of tutor-tuned neurons diminishes during development as the incidence of neurons that responds selectively to each bird’s own song increases, suggesting that tutor-tuned neurons are lost (Johnson and Bottjer, 1992, 1993, 1994) or re-tuned to provide a template of self-generated song (Volman, 1993; Zevin et al., 2004; Nick and Konishi, 2005a; Kojima and Doupe, 2007; Achiro and Bottjer, 2013). In accord with the latter idea, the emergence of selectivity for each bird’s own song is a ubiquitous signature of vocal learning across forebrain regions including HVC, LMAN, RA, basal ganglia, and thalamus (Margoliash, 1983; Margoliash and Konishi, 1985; Margoliash, 1986; Margoliash and Fortune, 1992; Volman, 1993; Doupe, 1997; Solis and Doupe, 1997; Person and Perkel, 2007).

Data based on playback of songs in anesthetized birds has highlighted the power of such experiments for studying mechanisms of vocal learning. However, several studies have shown that behavioral state is an important determinant of neural responsivity to song playback. Song-evoked responses can be elicited in sleeping as well as anesthetized zebra finches, and responses to song playback in adult birds tend to be highly similar between anesthetized and sleeping states (Dave et al., 1998; Dave and Margoliash, 2000; Nick and Konishi, 2001), encouraging the idea that anesthesia and sleep states are highly similar. In contrast, song-evoked responses are greatly diminished or absent in awake zebra finches (Schmidt and Konishi, 1998; Cardin and Schmidt, 2003; Rauske et al., 2003; Cardin and Schmidt, 2004b, a), which is reminiscent of the suppression of auditory responses to self-generated sounds in both vertebrate and invertebrate taxa (Suga and Shimozawa, 1974; Poulet and Hedwig, 2006, 2007; Eliades and Wang, 2008; Singla et al., 2017) (see Discussion). Here we report that responses to playback of different song types in both CORE and SHELL subregions of LMAN in sleeping juvenile birds are substantially different from those reported previously in urethane-anesthetized birds of the same age (Achiro and Bottjer, 2013). This difference is consistent with recent data showing that urethane anesthesia does not mimic sleep states (Mondino et al., 2021).

## Materials and Methods

### Subjects

All procedures were performed in accordance with the [Author University] animal care committee’s regulations. Five juvenile male zebra finches (*Taeniopygia guttata*) were used. Birds were bred in group aviaries and remained with their natural parents up until 33 days post-hatch (dph), at which time they and their father were removed from the main aviary and housed in an individual cage in the recording chamber in order to habituate them to the space. Experimental birds therefore received normal social-auditory experience and exposure to the tutor song (their father’s song) (Böhner, 1983, 1990; Mann et al., 1991; Mann and Slater, 1995; Roper and Zann, 2006).

### Electrophysiology

At 39 dph birds were anesthetized with isoflurane (1.5-1.8% inhalation) and placed in a stereotaxic apparatus. An electrode assembly consisting of eight tungsten-wire stereotrodes affixed to a movable microdrive was attached to the skull using dental cement such that the stereotrodes were implanted ^~^300 mm dorsal to LMAN CORE and SHELL. Each stereotrode was a twisted pair of polyester polyamide-imide overcoated tungsten wires (25 um diameter, California Fine Wire company, Grover Beach, CA) routed through fused silica capillary tubing (200 μm diameter). The assembly consisted of four posterior stereotrodes and four anterior stereotrodes; a silver wire, placed between the skull and skin, served as animal ground. Following surgery each bird was housed in a small individual cage in the recording chamber adjacent to the cage with the father; the father was removed 4-6 days later.

One to two days following surgery, the stereotrode assembly was connected to a recording headstage (HS-16, Neuralynx, Bozeman, MT) with a flexible cable connected to a commutator (PSR, Neuralynx); 15 channels of neural data were amplified, band passed between 300 and 5000 Hz (two Lynx-8 amplifiers, Neuralynx), and digitized at 32 kHz using Spike2 software (Power 1401 data acquisition interface, Cambridge Electronic Design). Audio and video were recorded coincident with neural activity: vocalizations were recorded to the 16^th^ channel using a lavalier microphone (Sanken COS-11D) mounted in the cage, and two USB-video cameras (30 FPS, ELP Day Night Vision, X000UPN1M5, HD 1080p) were placed at the front and side of the cage to record video files aligned to the neural activity. Two consecutive 60-min recordings were made between approximately 8 and 10 P.M. starting about one hour after lights off. Stereotrodes were manually advanced with the microdrive on consecutive days in the afternoon. The range of ages when recordings were made from LMAN CORE and/or SHELL ranged from 43 to 53 with a mean of 48.5 dph.

All birds received playback of four different songs: the bird’s own song (OWN, recorded within 24 hours prior to each recording), the bird’s tutor song (TUT), a juvenile conspecific song (JuvCon), and an adult conspecific song (AdlCon). The latter two songs served as control stimuli for OWN song and TUT song, respectively. JuvCon songs were age-matched to the age of the experimental bird’s OWN songs. The order of stimuli within a block of four songs was random without replacement, and the inter-stimulus interval was 30 ± 1 sec. Each song type was played back approximately 50 times at an amplitude of 56-59 dB, but only playbacks that occurred during sleeping periods were used for analysis (see below).

At the end of each experiment, birds were perfused (0.7% saline followed by 10% formalin), and brains were removed and postfixed before being cryo-protected (30% sucrose solution) and frozen-sectioned in the coronal plane (50 μm thick). Sections were Nissl stained with thionin to visualize stereotrode tracks and verify recording locations. The border between CORE and SHELL subregions of LMAN can be distinguished based on the density of magnocellular somata, which is low in SHELL relative to CORE.

### Data Analysis

A recording site was considered for analysis if it was confirmed histologically to be in either LMAN-CORE or LMAN-SHELL (excluding 50 μm on either side of the CORE/SHELL border). The evoked responses of LMAN neurons tend not to persist throughout song stimuli longer than 1 sec, as reported previously (Doupe, 1997; Solis and Doupe, 1997, 1999; Kojima and Doupe, 2007; Achiro and Bottjer, 2013). Therefore, response strengths calculated for song stimuli longer than 1 sec underestimate the actual response by averaging across both the early phasic response and the period of decreased response. To correct for this stimulus duration bias (e.g., longer songs underestimate true response strengths), all analyses were performed using neural data collected during the first second of song playback.

Periods of sleep were scored manually by two independent observers; as a conservative estimate, only periods ranked as sleep by both observers were used for analysis. Careful examination of the video files was used to mark sleeping periods as those in which birds were completely quiescent, displaying a regular pattern of deep rhythmic breathing with their eyes closed for at least 10 seconds. Sleeping periods were terminated at least 2 seconds prior to onset of large movements (e.g., wing movements) or eye-opening. As indicated above, only song playbacks that occurred during sleeping periods were included for analysis; the number of playbacks ranged from 16 to 48 (average 33 playbacks per song type in SHELL and 28 in CORE).

Movement artifact in multiunit neural recordings was correlated across recording channels and was eliminated or reduced using offline common average referencing: for each recording channel, the signal across the 14 remaining recording channels was averaged and subtracted from that channel to remove movement artifact (Ludwig et al., 2009). Noise was calculated as the standard deviation of the entire (2-hour) voltage recording, and minimum signal-to-noise ratio was set as three times the standard deviation; this threshold was used for spike detection. Single units were sorted from multiunit data by first automatically clustering units with KlustaKwik (KD Harris, University College London). KlustaKwik clusters were manually inspected across 18 different waveform features and further refined using MClust 4.4 (A. D. Redish, University of Minnesota). Clusters were included for analysis if < 1% of spikes had an interspike interval (ISI) < 2 ms.

We determined whether each single unit was responsive to song playback by testing for a significant change in firing rate (excitation or suppression) between baseline and each song type (Wilcoxon signed-rank test, p < 0.05). Baseline periods were defined as 1-sec periods immediately prior to stimulus playback, with the restriction that they must fall within sleep periods. For each song playback, the firing rates during the two closest baseline periods were averaged to generate a corresponding baseline value. To compare differences in firing rates across neurons, standardized response strengths (RS) were calculated as:

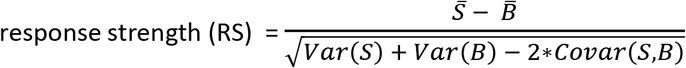

Where *S* is the average firing rate (spikes/sec) during stimulus, and *B* is the average firing rate during baseline, such that a positive value indicates an increased rate to a stimulus (excitation) and a negative value indicates a decreased rate (suppression).

To measure song selectivity for each song type for each cell, a difference score was calculated for “Song A” as follows: Song A_ΔRS_ = RS_SongA_ – RS_SongB_. For example, positive scores obtained by subtracting response strengths to OWN, AdlCon, and TUT (comparison songs) from JuvCon (reference song) would indicate selectivity for JuvCon. This measure is similar to the psychometric discriminability index d’ except that responses are standardized before being subtracted, as opposed to subtracting response strengths and then dividing by the standard deviations as in d’ (cf. Achiro & Bottjer 2013). Difference scores for song-suppressed responses were reversed in sign so that a positive difference score indicates a preference for the reference song over the comparison song while a negative difference score indicates a preference for the comparison song (see Results).

### Statistics

We used non-parametric statistics due to non-normal distributions of data, differing numbers of responses between song types, and differing numbers of neurons between CORE and SHELL regions. Differences in proportions were tested using χ^2^ tests, and differences in distributions were tested with Kolmogorov–Smirnov Z tests. Friedman tests were used to evaluate differences in RS between song types (as a repeated measure) within CORE and SHELL regions, whereas Kruskal-Wallis tests were used to evaluate differences between CORE and SHELL. Wilcoxon signed-rank tests were used to assess individual differences between song types, including Benjamini-Hochberg corrections for multiple comparisons (Benjamini and Hochberg, 1995). All values given as mean ± SEM.

## Results

### Different song types elicited different proportions of responses in both CORE and SHELL regions of LMAN

We recorded from CORE and SHELL subregions of LMAN in sleeping juvenile zebra finches (43 to 53 dph, mean = 48.5 dph). By this age juveniles have completed memorization of their tutor’s song and begun to practice their incipient song vocalizations. All neurons (n = 66, CORE; n = 104, SHELL) were tested with four different song types: each bird’s own song (OWN), each bird’s tutor song (TUT), an age-matched song from a juvenile conspecific (JuvCon) and an adult conspecific (AdlCon). Approximately half of the neurons in both CORE and SHELL showed a significant change in firing rate to at least one of the song types presented (CORE: 0.53, 35/66; SHELL: 0.47, 49/104); thus both regions showed similar levels of responsivity to song playback (χ^2^ = 0.57, p = 0.45). Proportions of significant playback responses varied by song type within both CORE and SHELL (CORE: χ^2^ = 13.8, p = 0.003; SHELL: χ^2^ = 12.6, p = 0.006) (Fig. 2 top panel; Table 1). JuvCon song elicited the highest proportion of responses whereas TUT evoked the lowest. Individual comparisons showed that the incidence of evoked responses to JuvCon was higher than that to TUT (CORE: p = 0.003; SHELL: p = 0.01, Fisher’s exact test, Benjamini-Hochberg corrected). JuvCon song also elicited a higher proportion of responses in LMAN-CORE neurons compared to AdlCon song (p = 0.04); no other comparisons between JuvCon and other song types were significant (p > 0.09 or higher).

**Figure 2.**
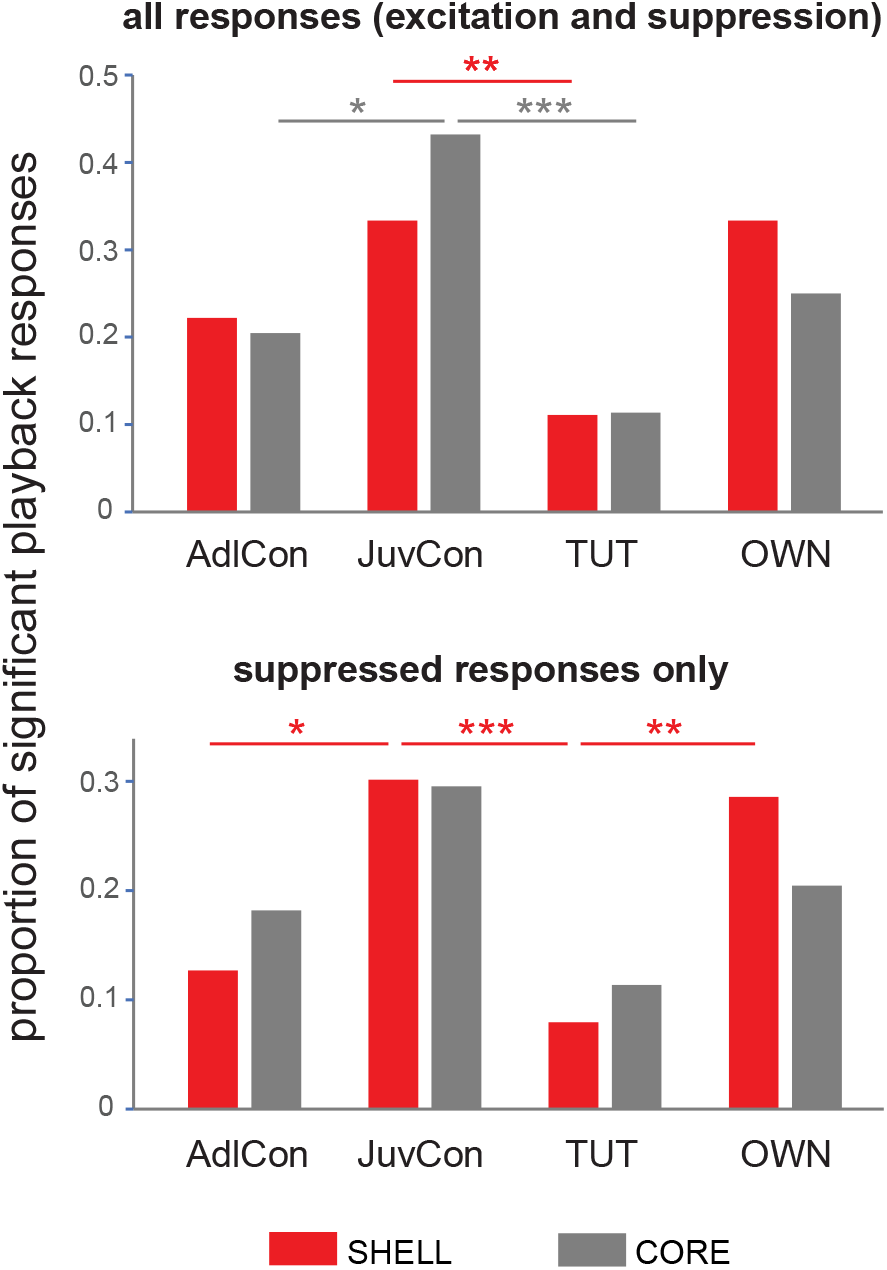
Proportion of significant responses to each song stimulus in CORE (gray) versus SHELL (red) neurons. Top: Proportions of excited and suppressed responses to playback of each song type (see Table 1). *p = 0.04, **p = 0.01, ***p = 0.003. Bottom: Proportions of suppressed responses to each song type. *p = 0.03, **p = 0.01, ***p = 0.006. AdlCon, Adult Conspecific song; JuvCon, Juvenile Conspecific song; TUT, tutor song; OWN, bird’s own song. n = 44 responses in 35 CORE neurons; n = 63 responses in 49 SHELL neurons.

**TABLE 1.**
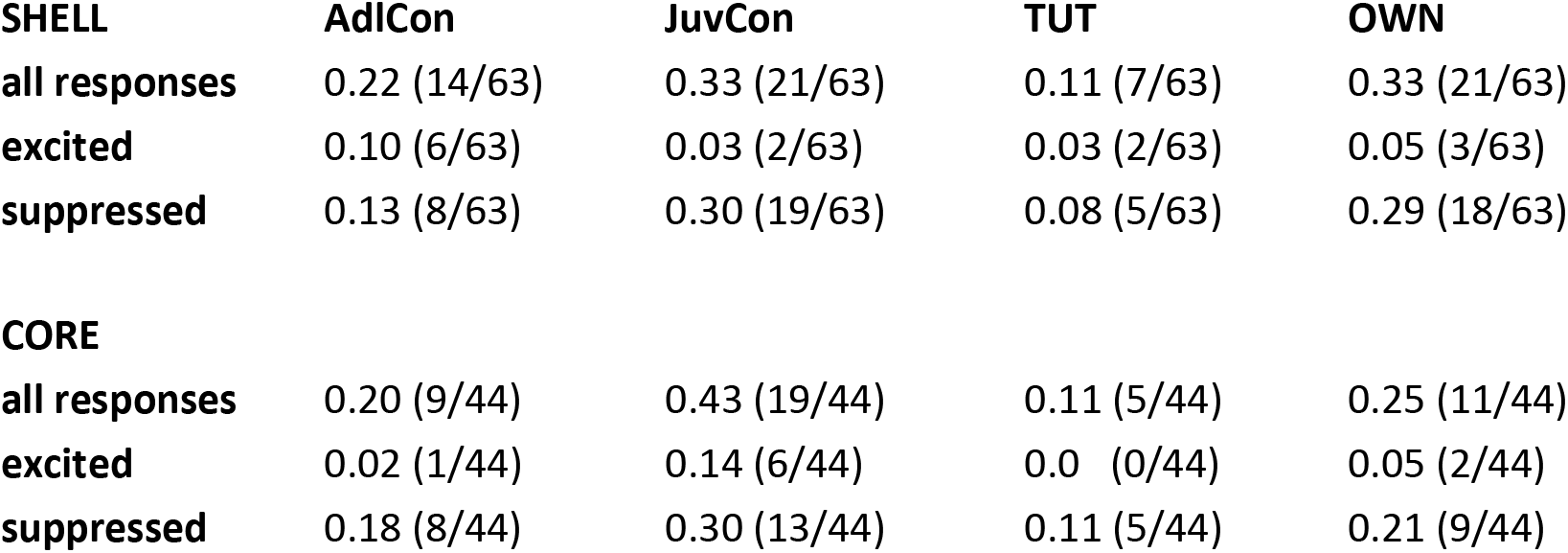
Proportions of significant responses by song type

| **SHELL** | **AdlCon** | **JuvCon** | **TUT** | **OWN** |
| --- | --- | --- | --- | --- |
| **all responses** | 0.22 (14/63) | 0.33 (21/63) | 0.11 (7/63) | 0.33 (21/63) |
| **excited** | 0.10 (6/63) | 0.03 (2/63) | 0.03 (2/63) | 0.05 (3/63) |
| **suppressed** | 0.13 (8/63) | 0.30 (19/63) | 0.08 (5/63) | 0.29 (18/63) |
| **CORE** |  |  |  |  |
| **all responses** | 0.20 (9/44) | 0.43 (19/44) | 0.11 (5/44) | 0.25 (11/44) |
| **excited** | 0.02 (1/44) | 0.14 (6/44) | 0.0 (0/44) | 0.05 (2/44) |
| **suppressed** | 0.18 (8/44) | 0.30 (13/44) | 0.11 (5/44) | 0.21 (9/44) |

Individual neurons were not broadly tuned: almost all neurons responded to either one or two of the four song types played; CORE neurons responded to 1.26 ± 0.07 different songs on average, whereas SHELL neurons responded to 1.29 ± 0.08. Figure 3 (left) shows that approximately 75% of neurons in both CORE and SHELL subregions responded to only one song type; the majority of the remaining cells responded to only two song types – no CORE neurons responded to three songs whereas 4% of SHELL neurons responded to three songs. The right side of Figure 3 depicts the song types to which each neuron responded, confirming that a low proportion of neurons in both CORE and SHELL responded to playback of TUT song, while relatively high proportions responded to both JuvCon and OWN songs.

**Figure 3.**
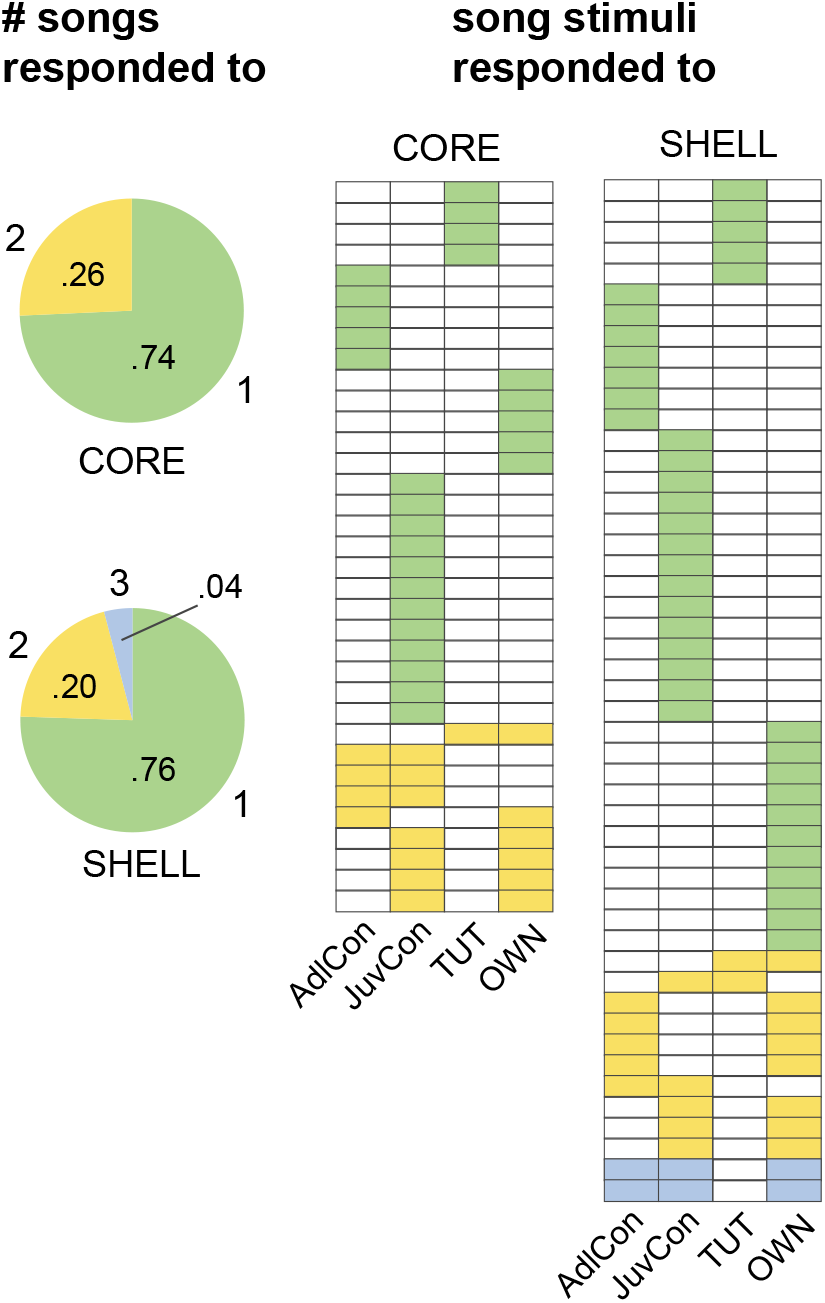
Single neurons were selectively tuned in both CORE and SHELL. Left: proportions of neurons that responded to different numbers of stimuli out of the four songs played; most neurons (^~^75%) responded to only one song stimulus in both CORRE and SHELL. Right: charts in which each row indicates the song stimuli to which each neuron responded (n = 44 responses in 35 CORE neurons; n = 63 responses in 49 SHELL neurons).

Although we found evidence of both excitation and suppression, the majority of cells within both CORE and SHELL showed only suppressed responses. Approximately 75% of cells in CORE and SHELL were suppressed by song playback, whereas relatively few cells responded with only excitation or a combination of excitation and suppression to different song types (Table 2). The dominance of suppressed responses was clear for all four song types, but was particularly pronounced for the two song types that elicited the highest percentage of responses – JuvCon and OWN (Table 1). We therefore examined the proportions of suppressed responses elicited by different song types (Fig. 2B). In contrast to comparison across all playback responses (Fig. 2 top), the bottom panel of Figure 2B shows that only SHELL neurons showed differential suppression between song types (SHELL: χ^2^ = 16.0, p = 0.001; CORE: χ^2^ = 4.99, p = 0.173). Within SHELL neurons, JuvCon evoked a higher incidence of suppressed responses compared to both TUT and AdlCon, but not OWN (TUT: p = 0.006; AdlCon: p = 0.034, Fisher’s exact test, Benjamini-Hochberg corrected). OWN song also evoked a higher proportion of suppressed responses compared to TUT (TUT: p = 0.011; AdlCon was marginally significant, p = 0.057; Fisher’s exact test, Benjamini-Hochberg corrected). Thus, the proportion of suppressed responses varied by song type in SHELL, but not CORE; within SHELL neurons both JuvCon and OWN elicited a high incidence of suppressed responses relative to AdlCon and (especially) TUT.

**TABLE 2.** Proportions of neurons by response type

| **SHELL** |  |  | **CORE** |  |  |
| --- | --- | --- | --- | --- | --- |
| (n=49) | # cells | proportion | (n=35) | # cells | proportion |
| excitation only | 8 | 0.163 |  | 4 | 0.114 |
| suppression only | 37 | 0.755 |  | 26 | 0.743 |
| both | 4 | 0.082 |  | 5 | 0.143 |

We are confident that we measured responses to song playback during periods of sleep since the use of behavioral criteria has been shown to be highly reliable (Szymczak et al., 1996; Low et al., 2008). We also noted that spontaneous firing rates (spikes/sec) during the night were higher during periods marked as waking compared to sleep periods (CORE waking 1.96, sleeping 1.51; SHELL waking 2.51, sleeping 2.07; Wilcoxon signed-ranks tests p < 0.0001 for both CORE and SHELL). To summarize data based on neuronal proportions in sleeping juvenile birds: 1) neurons in both subregions of LMAN responded in a selective fashion to song stimuli in sleeping birds during the period of early sensorimotor learning; 2) all individual songs were more likely to elicit suppression of firing rates rather than excitation, especially for JuvCon and OWN songs; 3) SHELL neurons showed a greater tendency towards suppressed responses to JuvCon and OWN songs compared to CORE neurons. Neurons at the population level evinced a preference for juvenile songs over adult songs, regardless of whether the juvenile song was self-generated (OWN) or produced by an age-matched conspecific bird (JuvCon).

This overall pattern of results contrasts markedly with that observed in a previous study in which birds of the same age were urethane-anesthetized rather than sleeping (Achiro and Bottjer, 2013). In that study, neurons in CORE were more likely to respond to playback compared to those in SHELL (0.89 versus 0.68), and neurons in both CORE and SHELL were much more likely to show excitation: CORE neurons never showed suppressed responses whereas ^~^80% of responses in SHELL neurons were excitatory and ^~^20% were suppressed. In addition, a large proportion of SHELL neurons exhibited a significant response only to TUT compared to those in CORE (0.28 versus 0.04), whereas a large proportion of CORE neurons responded to TUT plus other songs compared to SHELL neurons (0.43 versus 0.15) (Achiro and Bottjer, 2013). Approximately equal proportions of CORE and SHELL neurons showed responses only to OWN (CORE 0.15, SHELL 0.13). Thus SHELL neurons in anesthetized birds contain two distinct populations of neurons during early sensorimotor integration: one that responds only to the tutor song and a separate population that responds only to the bird’s own song. In general, CORE neurons in anesthetized birds are much more broadly tuned than SHELL neurons and show little evidence of responsivity to tutor song (see below and Achiro & Bottjer 2013). The current data did not replicate any of these patterns in sleeping birds (see Discussion and Table 3).

**TABLE 3.** Comparison of current results with those of Achiro & Bottjer 2013

|  | **Anesthetized** |  |  | **Sleeping** |  |
| --- | --- | --- | --- | --- | --- |
|  | CORE | SHELL |  | CORE | SHELL |
| % song-evoked neurons<sup>a</sup> | 89 | 68 |  | 53 | 47 |
| % suppressed neurons<sup>b</sup> | 0 | ~20 |  | 74 | 76 |
| % TUT-only responsive neurons<sup>c</sup> | 4 | 28 |  | 11 | 10 |
| selectivity score JuvCon versus TUT<sup>d</sup> | 0.13 | 0.43 |  | 0.47 | 0.45 |
| selectivity score JuvCon versus OWN<sup>d</sup> | 0.18 | 0.39 |  | 0.30 | 0.34 |

Anesthetized values are taken from Achiro & Bottjer (2013); sleeping values are taken from current study. The mean age of birds at which recordings were made in Achiro & Bottjer (2013) was 45.5 dph (range 43-47); the mean age of birds from recordings in the current study was 48.5 dph (range 43 to 53).
<sup>a</sup> percentage of neurons that responded to playback of at least one song.
<sup>b</sup> percentage of neurons that were suppressed by song playback (cells that showed suppression only are included for both studies).
<sup>c</sup> percentage of neurons that gave a significant response only to TUT and not to any other stimulus (out of 5 songs for Achiro & Bottjer, out of 4 songs for the current study).
<sup>d</sup> Selectivity scores for JuvCon versus TUT, OWN refer to average difference scores between standardized response strengths (see Methods). Scores from present study are for suppressed responses among JuvCon-selective cells, while scores from Achiro & Bottjer are for cells for TUT-selective and OWN-selective cells, respectively (Achiro & Bottjer scores include excited responses for CORE and excited and suppressed responses for SHELL); we chose to present scores for JuvCon-responsive cells from this study since so few neurons responded to TUT (Table 1).

### Response strengths were greater for songs that elicited a selective response

Figure 4 (top) shows absolute values of response strengths within all CORE and SHELL neurons for responses to each song type (including both excitatory and suppressed responses). This measure revealed no difference in firing rates between songs in SHELL but a significant difference in CORE (Friedman test: CORE, p = 0.039; SHELL, p = 0.257), reflecting a stronger response to JuvCon relative to other songs in CORE neurons. Given the relatively large proportion of significant responses to JuvCon song in both CORE and SHELL, we compared absolute values of response strengths across song types for the subset of neurons that responded significantly to JuvCon (n = 19 CORE, n = 21 SHELL). Figure 4 (middle) shows that this subpopulation in both CORE and SHELL exhibited a significantly higher firing rate to JuvCon compared to the other three song types (Friedman tests, p < 0.0001 in both CORE and SHELL; Wilcoxon signed-rank tests for JuvCon versus other song types always p < 0.004 or lower, Benjamini-Hochberg corrected). To determine whether this selective increase in firing rate was restricted to JuvCon-responsive neurons, we calculated firing rates for each subset of neurons that showed a significant response to the remaining three song types. A similar pattern was obtained for OWN-responsive, AdlCon-responsive, and TUT-responsive neurons, showing that single neurons that responded significantly to a given song type also showed a higher firing rate to that song type compared to other song stimuli. For example, OWN-responsive neurons in both CORE and SHELL had significantly higher response strengths to OWN compared to all other songs (Fig. 4, bottom; Wilcoxon signed-rank tests for OWN versus other song types in SHELL always p < 0.001; in CORE OWN versus AdlCon p = 0.005, versus JuvCon p = 0.003; versus TUT p = 0.001). (We did not perform statistical tests for TUT-responsive or AdlCon-responsive neurons due to relatively low n’s.)

**Figure 4.**
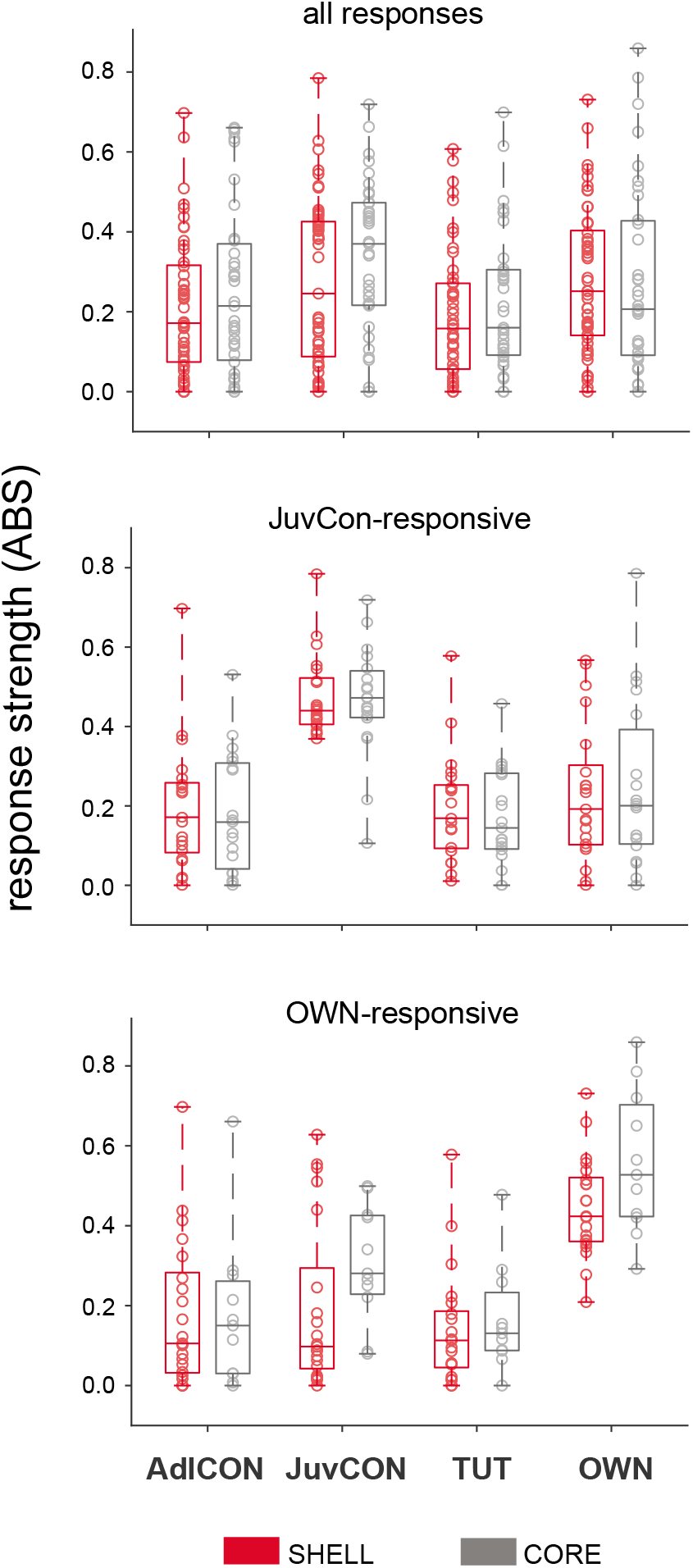
Standardized response strengths to each song stimulus across neurons. Box-and-whisker plots depict medians and first and third quartiles, whiskers indicate minimum and maximum values, and circles represent individual data points; all graphs depict absolute values (ABS) of standardized response strengths within CORE (gray) and SHELL (red) neurons. Top: response strengths to each song type (including both excitatory and suppressed responses, both significant and non-significant) for all CORE and SHELL neurons (n = 35 CORE, n = 49 SHELL). Middle: response strengths for the subset of CORE and SHELL neurons that showed a significant response to JuvCon song (n = 19 CORE, n = 21 SHELL). Bottom: response strengths for the subset of CORE and SHELL neurons that showed a significant response to OWN song (n = 11 CORE, n = 21 SHELL).

Given the prevalence of suppressed responses to JuvCon songs (Tables 1, 2), we examined neural selectivity between pairs of stimuli for JuvCon-suppressed neurons by calculating the difference in response strength between song types (see Methods). Response strengths to OWN, AdlCon, and TUT were subtracted from significantly suppressed JuvCon responses for each cell. A positive difference score indicates that a neuron preferred JuvCon song over comparison songs. Figure 5 shows cumulative distributions of difference scores in CORE versus SHELL neurons for JuvCon against each of the three other song types (n=13 CORE, n=19 SHELL). CORE and SHELL neurons clearly showed the same degree of preference for JuvCon song (Kolmogorov-Smirnov tests, p always > 0.87). A similar pattern of selectivity in CORE versus SHELL neurons was obtained when we compared cumulative distributions of difference scores for OWN (n=9 CORE, n=18 SHELL) against each of the three other song types (data not shown). Furthermore, very few neurons exhibited negative selectivity scores; the preponderance of positive scores in Figure 5 shows that cells that exhibited significant suppression to JuvCon almost never showed greater suppression to any other song stimulus. For example, only one SHELL neuron and no CORE neurons showed stronger suppression to AdlCon compared to JuvCon (Fig. 5). One-sample Wilcoxon signed-rank tests to assess whether the distributions were different from zero always yielded p values of < 0.0001.

**Figure 5.**
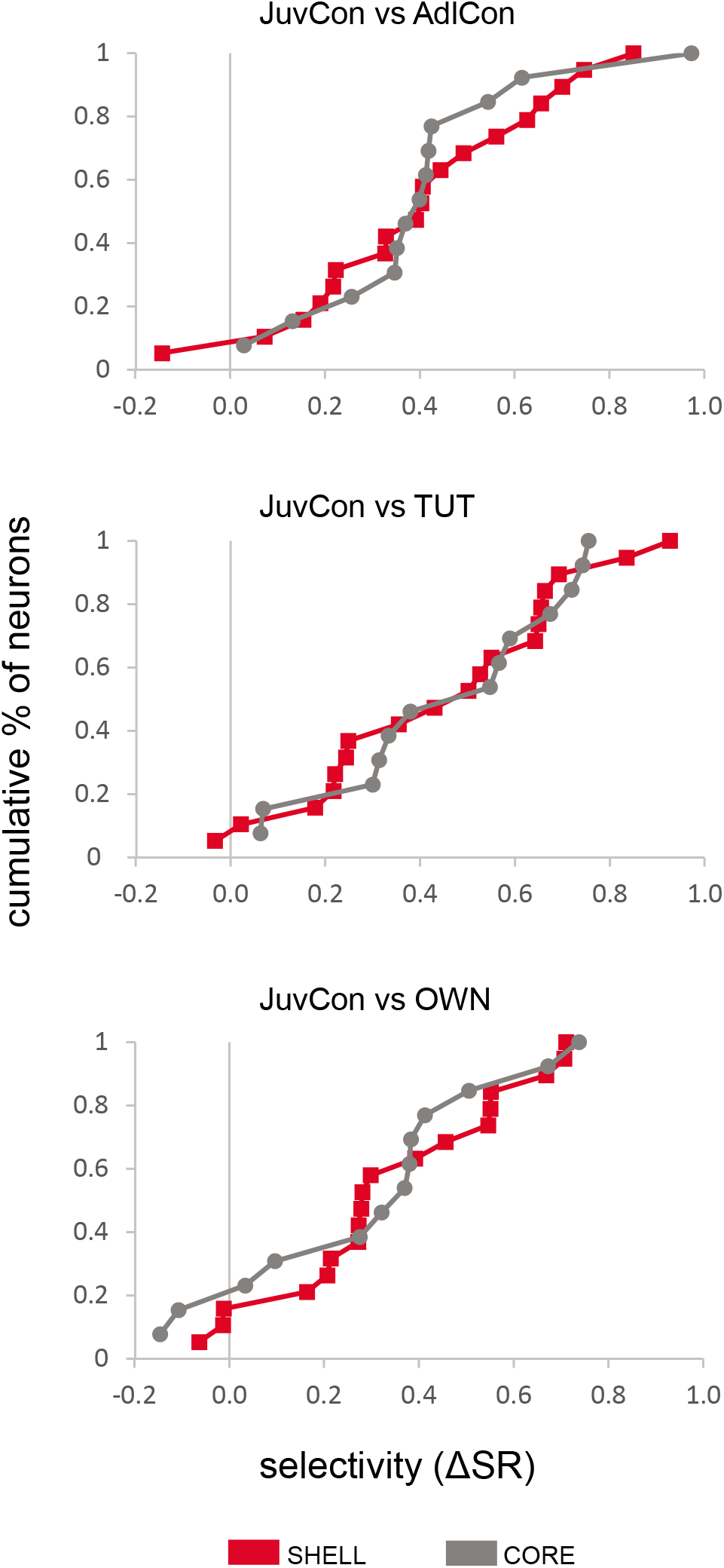
CORE and SHELL neurons were equally selective for JuvCon song. Each panel shows cumulative distribution functions of selectivity scores for JuvCon song compared to AdlCon (top), TUT, (middle) and OWN (bottom) (n = 13 CORE, n = 19 SHELL). Positive difference scores indicate a preference for JuvCon song over comparison songs, and show that both CORE (gray) and SHELL (red) neurons preferred JuvCon song over comparison songs to the same extent.

## Discussion

### Responses in awake versus sleeping or anesthetized states of adult animals

Many studies have shown that neural responses to song playback in the motor pathway of the song system, including the cortical regions HVC and RA (Fig. 1), are greatly diminished or absent in awake adult male songbirds but can be unmasked under anesthesia or in sleep (Dave et al., 1998; Schmidt and Konishi, 1998; Dave and Margoliash, 2000; Cardin and Schmidt, 2003; Rauske et al., 2003; Cardin and Schmidt, 2004b, a). This pattern may reflect, at least in part, a general tendency for responses in auditory and/or sensorimotor brain regions to be suppressed during self-generated sounds (Suga and Shimozawa, 1974; Poulet and Hedwig, 2006, 2007; Eliades and Wang, 2008; Singla et al., 2017). For example, neurons in auditory cortex of marmosets show suppression during vocal production; however, responses to their self-generated vocalizations are unmasked when auditory feedback is altered by realtime frequency shifts delivered through headphones (Eliades & Wang, 2008). One idea to arise from such findings is that learned signals from motor or other non-auditory inputs can predict auditory feedback and cancel responses to corresponding auditory sounds. A variant of this idea might explain the absence of song-evoked responses in awake songbirds; for example, motor circuits or pathways for efference copy might act to suppress auditory responses in an awake state even in the absence of active vocalizing. More broadly, the tendency for responses in awake or vocalizing animals to be suppressed is consistent with the idea that behavioral state can regulate a “gate” that controls auditory input.

In accord with this latter idea, the fact that responses to song playback in vocal-control regions of songbirds are absent in fully awake states but can be evoked under anesthesia or during sleep has been interpreted as changes in behavioral state, although no mechanism has been proposed as to the source of suppression in the awake state. In anesthetized and sleeping adult male songbirds, neurons throughout the song system are selectively tuned to each individual bird’s own song relative to conspecific songs or their OWN songs played in mirror-image reverse (Margoliash, 1983, 1986; Margoliash and Fortune, 1992; Vicario and Yohay, 1993; Nick and Konishi, 2001; Cardin and Schmidt, 2003; Person and Perkel, 2007). Cardin & Schmidt (2003) directly compared responses of HVC neurons in anesthetized, sleeping, and awake adult zebra finches; responses in both anesthetized and sleeping birds were consistently selective for OWN songs, whereas responses in waking birds were highly variable and not selective for OWN. Responses to playback in awake birds reflected the level of arousal: higher levels of arousal uniformly suppressed song-evoked responses in HVC (but had no effect in primary auditory cortex). The similarity of selective responses to OWN song in sleeping and anesthetized birds encouraged the idea that similar behavioral states underlie sleep and anesthesia.

### Responses in awake versus sleeping or anesthetized states of juvenile animals

Very few studies have examined responses to song playback in juvenile songbirds during sleep. Using behavioral criteria to assess sleep during song development, Nick & Konishi (2005b) reported that multiunit responses in HVC were strongest to tutor song in awake juvenile zebra finches during early sensorimotor integration, whereas OWN was preferred over tutor songs during sleep. Selectivity for OWN song changed over development in a pattern that tracked the current motor version of each bird’s song (Nick and Konishi, 2005a). Spontaneous patterns of spiking in HVC during sleep also change over song development: both firing rate and bursting increase with age (Crandall Nick et al. 2007).

EEG patterns are not a reliable indicator of sleep in juvenile zebra finches; the amplitude of 1-4 Hz activity (delta, an indicator of slow-wave sleep) did not vary between sleep and wake states in zebra finches between 45-65 dph (Nick and Konishi, 2005b; Crandall et al., 2007). This finding is consistent with the fact that the cortical EEG does not show evidence of state-dependent activity in early postnatal mammals (Gramsbergen, 1976; Frank and Heller, 1997; Blumberg et al., 2005). Even after EEG patterns differentiate (≥ 12 days postnatal in rodents), a long period of developmental changes ensues, which may be related to maturational changes that facilitate normal development of the nervous system (Khazipov and Luhmann, 2006; Cirelli and Tononi, 2015; Rensing et al., 2018).

Sleep is essential for vocal learning in juvenile zebra finches (Dave and Margoliash, 2000; Deregnaucourt et al., 2005; Crandall et al., 2007; Shank and Margoliash, 2009; Margoliash and Schmidt, 2010), which brings into question the influence of developmental changes during sleep in song-evoked activity, patterns of spontaneous spiking, and maturation of EEG patterns. Such changes within sensorimotor song regions may be related to substantial changes in the neural substrate for song learning (Alvarez-Buylla et al., 1988; Nordeen and Nordeen, 1988a; Nordeen and Nordeen, 1988b; Herrmann and Arnold, 1991; Johnson and Bottjer, 1992; Nordeen et al., 1992; Johnson and Bottjer, 1993, 1994; Livingston and Mooney, 1997; Foster and Bottjer, 1998; Iyengar et al., 1999; Kittelberger and Mooney, 1999; Livingston et al., 2000; Nixdorf-Bergweiler, 2001; Iyengar and Bottjer, 2002a, b; Bottjer, 2005; Miller-Sims and Bottjer, 2012; Garst-Orozco et al., 2014; Chung and Bottjer, 2021). For example, axonal projections that are present only during early stages of sensorimotor integration may mediate temporally restricted processes of song learning (Miller-Sims and Bottjer, 2012; Chung and Bottjer, 2021); in addition, refinement of axonal connectivity may represent either a morphological correlate of song learning or a necessary prerequisite for acquisition of song (Iyengar and Bottjer, 2002a, b). Developmental changes in sleep activity as well as in the neural substrate are likely to be related to changing patterns of responsivity to different song types at different stages of learning. A promising area for investigation lies in the extent to which developmental changes in sleep activity, the underlying neural substrate, selectivity for different song types, and maturation of vocal motor production are inter-related.

We are not aware of any previous studies that recorded the response of LMAN neurons to song playback during sleep in juvenile songbirds. Comparison of the present results in juvenile sleeping birds with those of Achiro and Bottjer (2013) in juvenile anesthetized birds clearly indicates that responses of LMAN neurons during sleep are substantially different from those recorded under urethane anesthesia in zebra finches during early sensorimotor integration. Salient differences in LMAN activity between this study and their previously published work are summarized in Table 3. Activity patterns in anesthetized birds revealed significant differences between CORE and SHELL for each of the measures listed in Table 3, whereas none of these measures varied between regions in sleeping birds. Two particularly striking differences are the dominance of suppressed responses in the present study, and the lack of a prominent neuronal subpopulation that responds selectively to tutor song in SHELL as is seen in anesthetized birds. These differences raise the question of when and how the tutor-tuned SHELL neurons are utilized in the service of learning. Perhaps our sleep conditions were not conducive to eliciting responses from tutor-selective neurons, in which case they may have an important sleep-related function under more normal conditions (e.g., in sleeping birds that are not wearing an electrode array attached to a cable). Or perhaps these neurons are actively involved in some aspect of learning during sleep but are gated off from activation via auditory playback. Another possibility is that tutor-tuned neurons can be activated during awake states (as for HVC neurons of juvenile birds, Nick & Konishi, 2005), particularly during singing. If tutor-tuned neurons are activated only during singing in awake states, their activity might be difficult to identify in the context of motor-related activity (Achiro et al., 2017).

### Comparing urethane anesthesia and different sleep states

Prior to beginning our study we assumed that responses to song playback during sleep in LMAN of juvenile birds would replicate results in anesthetized birds (which would have enabled direct comparison of playback-evoked patterns as in anesthetized birds and singing activity in awake birds). Because we did not intend to study sleep-related factors we made no effort to characterize different stages of sleep in relation to playback. Despite the fact that EEG patterns do not correlate with sleep stages in juvenile animals (Gramsbergen, 1976; Frank and Heller, 1997; Blumberg et al., 2005; Nick and Konishi, 2005b; Crandall et al., 2007; Cirelli and Tononi, 2015), different states of sleep and/or ultradian rhythms may nevertheless influence song responsivity. If so, different sleep states might provide a possible alternative explanation of the stark differences we observed between song-evoked activity in LMAN of sleeping versus anesthetized juvenile zebra finches. Robust responses to song playback are observed during slow wave sleep in HVC of adult zebra finches (Nick and Konishi, 2001). We are not aware of any studies that have compared song-evoked responses during REM (rapid eye movement) versus non-REM sleep. It would be interesting to correlate responsivity to song playback with EEG patterns in adult birds, taking into account that episodes of different sleep states are quite brief (less than 30 sec in adult budgies) and slow wave sleep decreases through the night while REM sleep increases (Canavan and Margoliash, 2020). It is not clear how informative this approach might be in young songbirds given that EEG patterns are not a reliable indicator of sleep states in juvenile animals, although it is nevertheless possible that a given song type could elicit different neural responses in sensorimotor song regions depending on EEG activity.

The similarity of selective responses to OWN songs under sleep and anesthesia in HVC neurons of adult songbirds has encouraged the idea that behavioral states are highly similar between the two conditions. However, this idea has not been extensively tested in either birds or mammals. Some studies have suggested that urethane anesthesia mimics sleep, based on alternation of EEG patterns between a slow-wave state that resembles non-REM sleep and an “activated” state with features of both REM sleep and waking (Clement et al., 2008; Pagliardini et al., 2013; Tisdale et al., 2018). Recent work has not supported this idea, based on detailed comparisons that measure several correlates to define conscious (waking) versus unconscious (sleeping) states, including power spectra of EEGs, synchronization between high-frequency (gamma) oscillations in different brain regions, directional patterns of activation, and temporal complexity of neural oscillations (Mashour and Hudetz, 2018; Kelz and Mashour, 2019; Mashour et al., 2020). A recent study that performed within-subject comparisons of sleep versus urethane anesthesia in rats reported that these EEG correlates of consciousness were significantly lower during anesthesia compared to sleep (Mondino et al., 2021). For example, normalized power of delta oscillations was higher during both “REM-like” and “non-REM-like” states of urethane anesthesia compared to their respective REM and non-REM states during sleep. In addition, despite qualitative similarities between REM and non-REM EEG patterns in sleep versus urethane anesthesia, clear-cut differences in temporal complexity existed between them. These authors concluded that urethane induces a pattern of “sustained unconsciousness” dissimilar from that of sleep. Thus, it seems likely that differences in patterns of brain activity between sleep and anesthesia could underlie the different responses to song playback that we observed in LMAN of juvenile zebra finches between the current study and previous work by Achiro and Bottjer (2013). If so, that would suggest that urethane anesthesia is more effective at removing one or more gates of song-evoked activity compared to sleep.

